# Spatial multiomic profiling reveals distinct fibrotic epithelial niches in idiopathic pulmonary fibrosis

**DOI:** 10.1101/2024.12.10.627749

**Authors:** Bin Liu, Konain Bhatti, Iain D. Stewart, James May, Maria Concetta Zarcone, Nathalie Lambie, Nik Matthews, Rachel L Clifford, Charlotte H. Dean, Peter M. George, Richard Hewitt, Elena Lopez-Jimenez, Maddy Parsons, Simon R Johnson, Anna K. Reed, Elisabetta A. Renzoni, Jody Rosenblatt, Amanda L. Tatler, Isabel Uwagboe, Louise V Wain, Athol U. Wells, R. Gisli Jenkins, Alison E. John

**Author notes:** Correspondence to R Gisli Jenkins: Margaret Turner Warwick Centre for Fibrosing Lung Disease, National Heart and Lung Institute, Imperial College London, Guy Scadding Building, Cale Street, London, SW3 6LY.

## Abstract

**Background:** Idiopathic pulmonary fibrosis (IPF) is a progressive, fatal disease characterised by excessive extracellular matrix deposition within the lung. Recent advances in single-cell RNA sequencing have identified distinct fibrotic populations, yet their origins and spatial relationships remain incompletely understood.

**Methods:** Using spatial transcriptomics and Hyperion imaging mass cytometry we compared the cellular composition in formalin fixed paraffin embedded fibrotic lesions (n=9 patients) with control lung (n=9), and cellular interactions were inferred using CellChat V2 ligand-receptor analysis. Monolayers of airway epithelial cells were used to identify changes in keratin (KRT) expression following cell detachment and cyclical mechanical stretch.

**Results:** Spatial multiomics profiling of human lung cells confirmed the *in situ* localisation of previously described IPF-enriched populations, and identified a previously unrecognized KRT5^low^/KRT17⁺ epithelial population derived from airway basal cells that progressively acquiring molecular features of aberrant basaloid cells, forming a unique fibrotic niche enriched with the Secreted Phosphoprotein 1 (SPP1) positive macrophages. Functional studies demonstrated that epithelial detachment and cyclical mechanical stretch drive KRT5 reduction, providing a mechanism for the emergence of this transitional state. In addition, we also identified distinct immune–stromal niches enriched in lymphocytes and alveolar fibroblasts.

**Conclusion:** These findings delineate distinct fibrotic epithelial niches in IPF and support a model in which epithelial loss induces aberrant basaloid differentiation and fibroblast activation, with subsequent airway traction and epithelial detachment generating a secondary niche enriched in basal-derived KRT5^low^/KRT17 cells and SPP1⁺ macrophages.

## Introduction

Idiopathic pulmonary fibrosis (IPF) is a chronic, progressive, and fatal interstitial lung disease characterized by matrix deposition within the lung which leads to a relentless decline in pulmonary function. IPF is more common with increasing age, and incidence rates are rising with an estimated 1-13 cases per 100,000 population globally(1). IPF is due to the accumulation of environmental risk factors in people with a genetic predisposition. Despite advances in understanding of pathogenesis and improved treatment options, IPF remains a condition with high morbidity and mortality with median survival rates from 3 to 5 years after diagnosis(2, 3).

Recent advances in single-cell sequencing have accelerated understanding of the diverse cellular populations contributing to IPF pathogenesis. A distinct population of Keratin 5 negative (KRT5^-^) but Keratin 17 positive (KRT17^+^) epithelial cells has been identified in fibrotic regions of the lung. These KRT5^-^/KRT17^+^ cells also express markers such as Vimentin, matrix metalloproteinase (MMP7), tumour protein p63 (TP63), and SRY-box2 (SOX2), indicating their progenitor-like nature(4–7). Similarly, a unique population of collagen-producing fibroblasts characterized by increased production of type 1 collagen and the expression of collagen triple helix repeat containing 1 (CTHRC1+) have been identified(8, 9). Finally, macrophages that express Secreted Phosphoprotein 1 expressing macrophages (SPP1+)(10) and secrete osteopontin which promotes extracellular matrix production and trigger myofibroblasts activation(11), have been detected.

Spatial multi-omics provide powerful tools for understanding cellular heterogeneity by combining high-dimensional molecular profiling with spatial context. Spatial transcriptomics permits gene expression mapping within intact tissue sections by revealing the spatial distribution of mRNA transcripts, helping to elucidate cellular interactions and microenvironmental influences(12, 13). Furthermore, high-multiplex tissue imaging enables the simultaneous imaging of multiple proteins at the single-cell level *in situ*, providing further detailed information about cellular phenotypes and states while preserving spatial information(14). Combining both technologies enables a deep molecular phenotyping of the cellular heterogeneity in IPF and how these cells interact to contribute to the fibrotic niche.

In this study, spatial transcriptomics was combined with Hyperion imaging mass cytometry to comprehensively assess the fibrotic niche within human lung tissue obtained from patients with IPF. We observed a profound reduction in AT1 cells in fibrotic lung, along with an increase in both KRT5^+^/KRT17^+^ basal, KRT5^-^/KRT17^+^ aberrant basaloid cells and notably, a hitherto undescribed KRT5^low^/KRT17^+^ transitional basal cell population. The KRT5^low^/KRT17^+^ cells communicates primarily with SPP1+ macrophages, yet share gene expression signature with AT2-derived aberrant basaloid cells, which predominantly interact with activated myofibroblasts. In vitro analysis of airway basal epithelial cells revealed that monolayer detachment and cyclical stretch leads to loss of KRT5. These data highlight a novel and important pathway where airway derived progenitor cells contribute to aberrant alveolar repair in IPF, promoting a unique airway derived fibrotic niche.

## Materials and Methods

### Human tissue collection and tissue microarray construction

IPF lung tissues were sourced from the Nottingham NIHR Biomedical Research Centre under ethical approval (REC 08/H0407/1). Non-IPF tissues were collected from Clinical Research Facility (CRF), Royal Brompton Hospital under the ethical approval (NRES 20/SC/0142). Lung tissues were formalin-fixed and paraffin-embedded prior to tissue microarray (TMA) construction. 1 mm cores were sampled from IPF (4 cores) and control (2 cores) tissues.

### CosMx SMI data processing and analysis

Machine learning-driven single cell segmentation was performed using AtoMx Spatial Informatics Platform (SIP)(15, 16). For data quality control (QC), cells with negative probe greater than 0.5 and or features less than 20 were filtered out. In total, 180,067 cells were profiled, and raw counts of all cells were exported as a Seurat object from AtoMx SIP for downstream analysis. Total counts normalization was applied. Following normalization, principal component analysis (PCA) and then Uniform Manifold Approximation and Projection (UMAP) and t-distributed Stochastic Neighbor Embedding (tSNE) were performed for dimension reduction and visualisation using Seurat v5(17).

### Cell type annotation

We applied a hierarchical stratergy for cell type annotation, starting with Leiden clustering and aggregating clusters into the main cell lineages based on Human Lung Cell Atlas (HLCA) v2(18) level1 markers which are included in the CosMx 1000-plex gene panel: Immune markers: (CD53, PTPRC, COTL1, CXCR4, FCER1G, SRGN, CD52); Epithelial markers: (KRT7, PIGR, KRT8, KRT19, TSPAN8, CSTD2, CXCL17); Endothelial markers: (PECAM1, VWF, RAMP2, IGFBP7, CLEC14A); and Stromal markers: (TPM2, DCN, MGP, CALD1, LUM, TAGLN, COL1A1). These four lineage subsets were then used for downstream cell subtype annotation using the Insitutype algorithm(19), applying semi-supervised cell typing.

### Inference of inter-niche and inter-cell communication using CellChat V2

For inference of cell communication, we first constructed a Niche assay from CosMx spatial transcriptomics data by aggregating local cell-type composition and grouping similar neighbourhoods into recurrent niches. Clustering of neighbourhoods revealed seven distinct niches (N1–N7). N1: MyoFB niche, N2: Airway niche,N3: KRT5-/KRT17+ niche,N4: Immune niche,N5: Vascular niche,N6: Alveolar niche, N7: SPP1+ macrophage niche. CellChat V2(20) was used to infer spatially proximal inter-niche communication and intra-niche intercellular communication. CellChat DB V2 was used as the ligand-receptor (L-R) interaction database. A total of 358 L-R pairs were extracted from the NanoString 1000-plex RNA probes panel and used for further analysis.

### Hyperion IMC data processing and analysis

Preprocessing, segmentation, and feature extraction of multi-channel Hyperion IMC images were achieved using the IMC Segmentation Pipeline, as previously described(21). Image based single cell spatial analysis of the IMC data was performed using imcRtools and cytomapper R package.

### Imaging Flow Cytometry

Imaging flow cytometry of KRT5 in ihBECs was performed using the ImageStream MK II system. Cells were permeabilised and fixed using eBioscience™ Foxp3 / Transcription Factor Staining Buffer Set (Thermo Fisher Scientific) following the supplier’s protocol. Following fixation, cells were stained for KRT5 (2ug/mL) antibody for 30mins at 4° in the dark. Gating and downstream analysis was performed using IDEAS software.

All additional details of the materials and methods section are provided in an online data supplement.

## Results

### Spatial analysis of control and IPF human lung tissues identifies loss of normal alveolar cells and enriched fibrotic cellular populations

A tissue microarray constructed from nine IPF patients and nine controls were assessed by spatial transcriptomics, yielding 180,067 cells (141,070 IPF and 38,997 controls), with an average of 110 transcripts per cell analysed. Dimensionality reduction revealed four major cellular compartments corresponding to epithelial, stromal, endothelial, and immune lineages (Figure 1A). Spatial mapping of these transcriptionally defined compartments to tissue sections confirmed expected anatomical localization. (Figure 1B).

**Figure 1.**
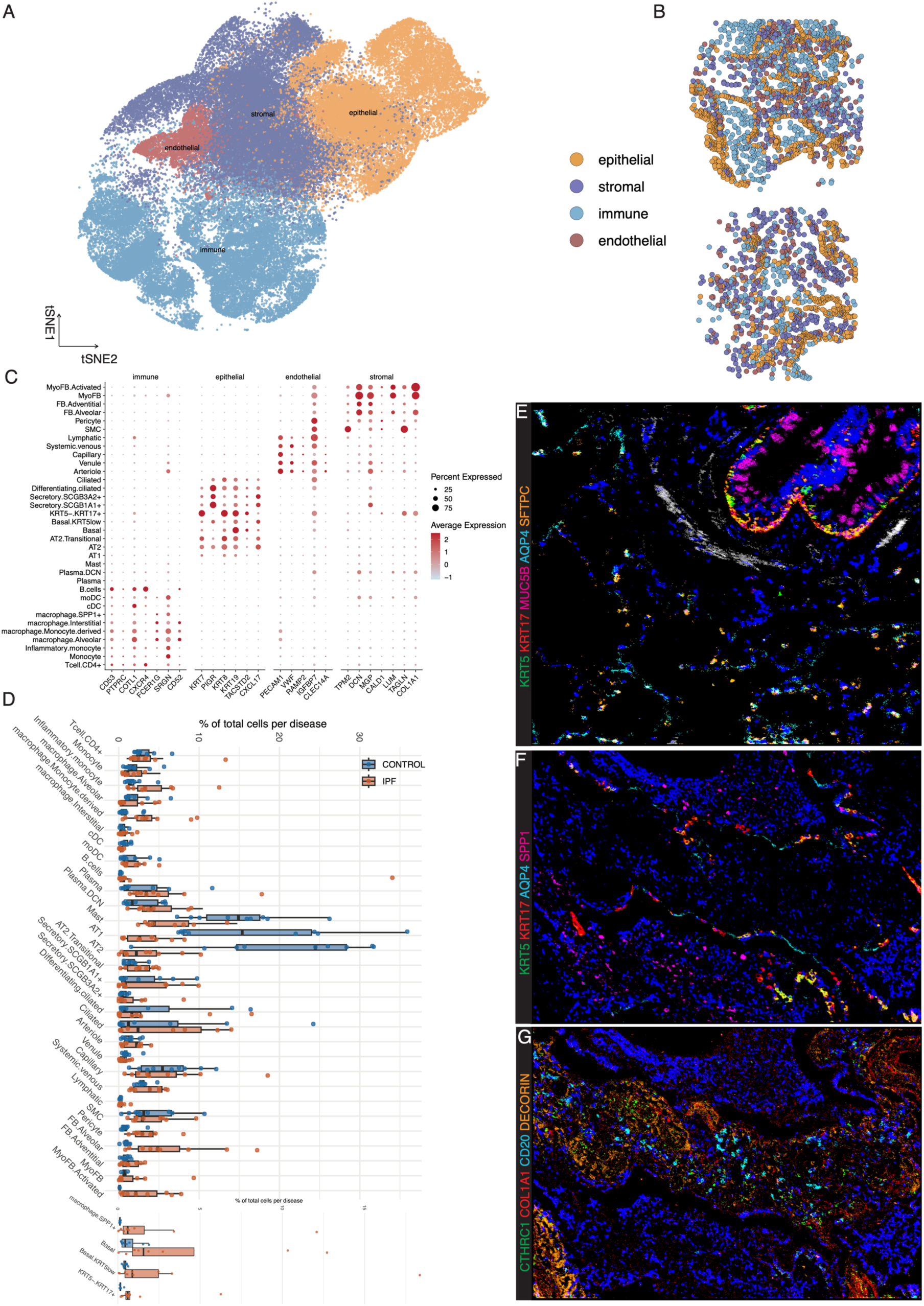
Cellular composition of control and idiopathic pulmonary fibrosis (IPF) lungs profiled by CosMx Spatial Molecular Imaging. **A** t-SNE embedding of all CosMx single-cell transcriptomic profiles, coloured by major lineage (epithelial, immune, endothelial, stromal). B Representative CosMx images showing the spatial distribution of epithelial, immune, endothelial, and stromal lineages in lung tissue sections. C Dot plot showing expression of HLCA v2 level 1 marker genes across the final 34 annotated cell types. Dot size represents the proportion of cells expressing each gene and colour intensity indicates average expression, confirming annotations within epithelial, immune, endothelial, and stromal lineages. D Boxplots showing the proportion of each cell type, expressed as a percentage of total cells per sample, in control (n = 9) and IPF (n = 9) lungs. Points represent individual samples. Boxplots indicate the median, interquartile range (box), and whisker extension. Colours correspond to disease state. E-G representative Hyperion imaging mass cytometry images of E Control lung tissue showing AQP4⁺/SFTPC⁺ alveolar parenchyma and KRT5⁺/KRT17⁺/MUC5B⁺ airway epithelium surrounded by αSMA⁺ regions. F IPF lung tissue with KRT5⁻/KRT17⁺ epithelial cells located in close proximity to KRT5+/KRT17⁺ basal cells, AQP4⁺ AT1 cells, and near SPP1⁺ macrophages G IPF lung tissue showing CD20⁺ B cells positioned near CTHRC1⁺ fibroblasts within collagen I– and decorin-rich fibrotic region. AT1, alveolar type 1; AT2, alveolar type 2; FB, fibroblast; MyoFB, myofibroblast; SMC, smooth muscle cell; DCN, decorin; moDC, monocyte-derived dendritic cell; cDC, classical dendritic cell; NK, natural killer; Mast, mast cell

To generate high-resolution cell type classification, we applied a semi-supervised annotation approach using the Insitutype algorithm, guided by an expression matrix derived from the publicly available single-cell RNA-seq dataset (GSE227136). This resulted in 34 discrete subtypes that clustered into the four major cell classes based on HLCA v2 level 1 marker gene expression (Figure 1C). Analysis of cell type proportions revealed disease-associated alterations. Specifically, alveolar type 1 and type 2 (AT1 and AT2 cells) cell populations were significantly reduced in IPF, whereas basal (KRT5^+^/KR17^+^, KRT5^low^/KRT17^+^), and aberrant basaloid (KRT5^−^/KRT17^+^) epithelial cells were all substantially expanded (Figure 1D). Within the stromal compartment, a marked increase in alveolar fibroblasts and activated myofibroblasts was observed in IPF compared with controls. In addition, immune populations were also enriched, including monocyte derived and SPP1^+^ macrophages as well as decorin expressing plasma cells (Figure 1D).

To validate the findings from transcriptomic data, we applied multiplex imaging mass cytometry on adjacent sections to assess protein-level changes. Unsupervised clustering following single-cell segmentation revealed 15 distinct cell populations, including IPF-enriched cell populations (Figure E2A, B). Eighteen markers overlapped between CosMx SMI and Hyperion IMC marker panel, and most cells expressed at least one of these shared markers in both datasets (Figure E2C, D). In control tissues, KRT17^+^/KRT5^+^ basal cells were confined to the airways adjacent to secretory airway epithelial cells (MUC5B^+^) and absent from the alveolar parenchyma where high numbers of cells expressing aquaporin-4 (AQP4)⁺ and surfactant protein C (SFTPC)^+^ were observed (Figure 1E). In contrast, IPF-enriched populations displayed distinct spatial distributions: KRT5⁻/KRT17⁺ epithelial cells lined fibrotic regions and were positioned near KRT5⁺/KRT17⁺ basal cells (Figure 1F, G). Additionally, KRT5⁻/KRT17⁺ cells were observed adjacent to AQP4⁺ AT1 cells and in close proximity to SPP1^+^ macrophages (Figure 1F). Meanwhile, CTHRC1^+^ fibroblasts were primarily located within the fibrotic stroma, exhibiting high levels of collagen I expression (Figure 1G). CD20+ lymphocytes were also present within these matrix-rich fibrotic regions, in proximity to CTHRC1^+^ fibroblasts (Figure 1G).

### Profiling spatial cellular neighbourhoods reveals distinct fibrotic niches

To investigate spatial interactions and tissue-restricted niches, cell neighbourhood analysis was performed as previously described(17, 21) using the CosMx spatial transcriptomic dataset. Seven niches with distinct cellular compositions were identified in CosMx dataset (Figure 2A). *In situ* projection and quantification of cell centroids revealed that control lungs were dominant by an alveolar niche (N6 consisting mainly of alveolar cells), an airway niche (N2 enriched for airway epithelial cells), and a vascular niche (N5, defined by smooth muscle and endothelial cells). In contrast, IPF lungs demonstrated a marked reduction of N6, an expansion of N2, and the emergence of abnormal niches characterised by disease-associated cell populations (Figure2A, C). These included four distinct fibrotic niches: (N1) consisting of activated myofibroblasts-and immune cells; (N3) KRT5-/KRT17+ aberrant basaloid cells, activated myofibroblasts and alveolar macrophages; (N4) an immune-stromal cell niche driven by lymphocytes and fibroblasts; and (N7) an SPP1+ macrophages predominant niche (N7) (Figure2A-2C). To further explore potential communication between niches, we applied CellChat to infer niche-niche interactions. Circle plots scaled by interaction weight revealed sparse interactions in control tissue, whereas IPF exhibited numerous and stronger interactions, particularly involving N3 and N6 (Figure 2D. Ligand-receptor(LR) pair aggregation based on pathway revealed dominant SPP1, collagen, and FN1 signalling in IPF niches (Figure 2E), highlighting LR pairs such as SPP1-CD44 and COL6A-CD44 across multiple fibrotic niches (Figure E3).

**Figure 2.**
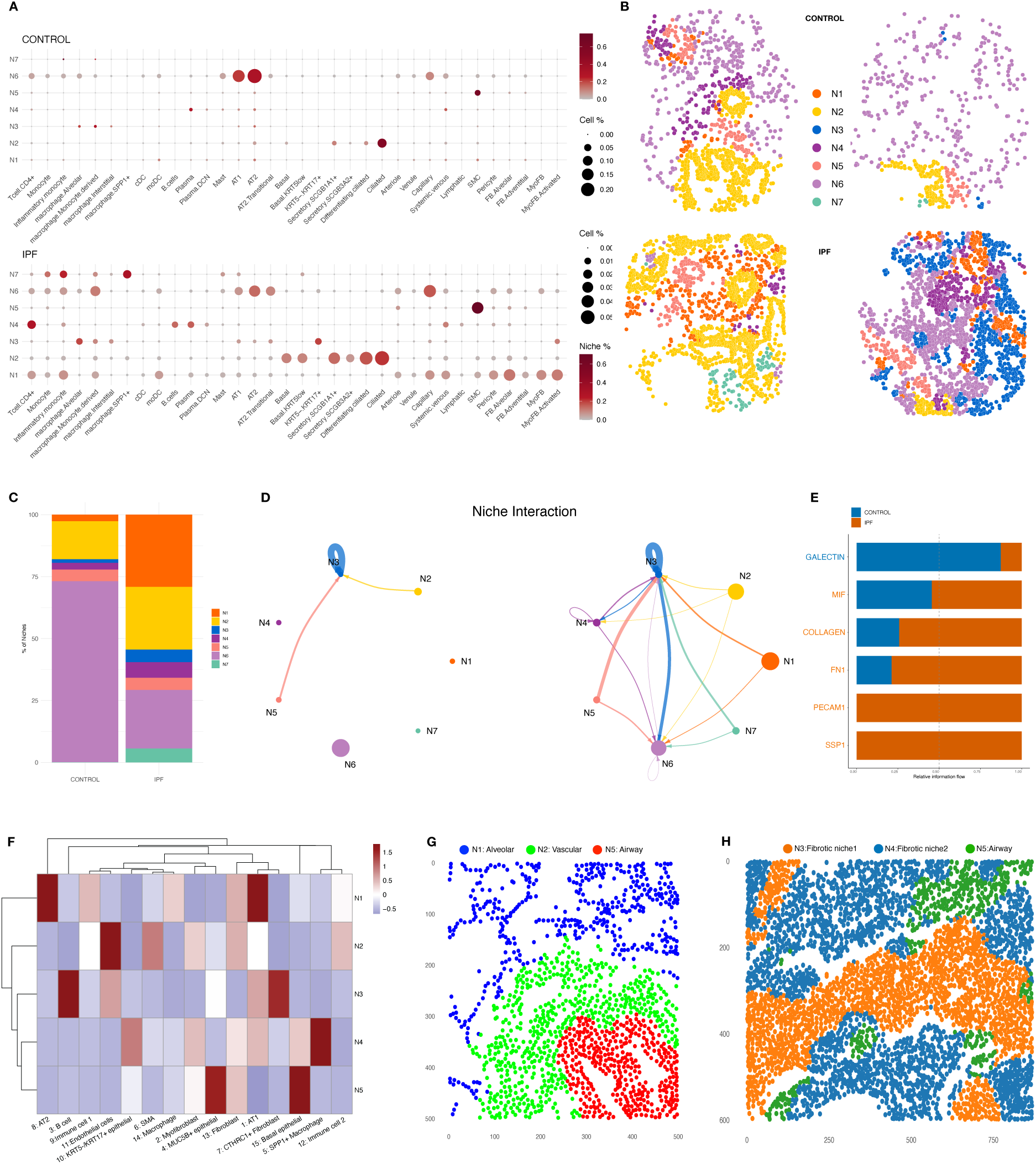
Spatial niche architecture in control and IPF lungs. CosMx analysis of spatial niches in control and IPF lungs **(A,B)**. **A** Dot plot showing the cellular composition of niches (N1–N7) in control (top) and IPF (bottom). Dot size represents the proportion of each cell type relative to the total cell number, while dot colour indicates the proportion of each cell type within a given niche. **B** Spatial maps of cell centroids coloured by niche identity (N1–N7) in control (top) and IPF (bottom). N1 active myofibroblast niche, N2 airway niche, N3 KRT5⁻/KRT17⁺ niche, N4 immune niche, N5 vascular niche, N6 alveolar niche, and N7 SPP1⁺ macrophage niche. **C** niche composition of control and IPF tissue shown as a stacked bar plot. Cellchat analysis of inferred ligand–receptor interactions depicting **D** the inter-niche communication in control and IPF lung tissues. Nodes represent cell types and edges indicate predicted interactions, with edge width proportional to interaction strength, and **E** top ranked pathway aggregated by ligand-receptor pairs in control and IPF. Hyperion IMC analysis of spatial niches in control and IPF lungs **(F–H)**. **F** Heatmap showing the cell type composition of niches (N1–N5). Rows correspond to niches and columns to cell types. Colours indicate relative enrichment of each cell type across niches (dark red, enriched; dark blue, depleted; white, average). N1 alveolar niche, N2 vascular niche, N3 fibrotic niche 1, N4 fibrotic niche 2, and N5 airway niche. D, E Representative spatial maps of cell centroids coloured by niche identity (N1–N5) in control **(G)** and IPF **(H)**.

In parallel, cell community analysis of Hyperion IMC data revealed five niches (Figure 2F), including an alveolar niche enriched for SFTPC+ AT2 and AQP4+ AT1 cells, an airway niche composed primarily of basal (KRT5^+^/KRT17^+^) and goblet (MUC5B^+^) epithelial cells, and a vacular niche dominated by αSMA^+^ cells (Figure 2G). Hyperion IMC further identified two distinct fibrotic niches: the first fibrotic niche predominantly featured SPP1^+^ macrophages, KRT5^-^/KRT17^+^ epithelial cells, and basal epithelial cells (Figure1E, Figure 2F, 2H), while second fibrotic niche was defined by fibrotic fibroblasts (CTHRC1^+^/COL1^+^) in association with CD20+ lymphocytes (Figure1G, Figure 2F, 2H).

### Distinct origins of fibrotic epithlium contribute to fibrotic niches

To further understand the nature and origin of aberrant epithelial cells, we analysed all epithelial cells separately within fibrotic lung. UMAP visualisation revealed two major epithelial populations corresponding to alveolar and airway lineages, including a previously undescribed KRT5^low^/KRT17^+^ cell, along with a small convergent population of KRT5^-^/KRT17^+^ cells (Figure 3A). Trajectory inference suggested that KRT5^-^/KRT17^+^ cells could arise from AT2 cells consistent with prior reports (Figure 3B), but also indicated a possible origin from airway basal cells (Figure 3C). To further resolve the relationship between basal, KRT5^low^/KRT17^+^ and KRT5^-^/KRT17^+^ epithelial cells, we examined pseudotime expression dynamics of aberrant basaloid markers. KRT5^low^/KRT17^+^ cells occupied an intermediate state, with elevated expression of growth differentiation factor 15 (GDF15), matrix metallopeptidase 7 (MMP7), transmembrane 4 L six family member 1 (TM4SF1), SRY-box transcription factor 4 (SOX4), integrin subunit beta 6 (ITGB6), integrin subunit alpha V (ITGAV), and cyclin-dependent kinase inhibitor 1A (CDKN1A) relative to basal cells, but lower than KRT5^-^/KRT17^+^ cells (Figure3D). Spatial mapping centroids (Figure3G, 3J) of these KRT5^low^/KRT17^+^ cells revealed enrichment in fibrotic epithelial (PanCK+) regions (Figure3E-3F, 3H-I), often interspersed with AT2 cells and KRT5^-^/KRT17^+^ cells (Figure 3J). Distance analysis demonstrated co-localisation between AT1, AT2, AT2 transitional and KRT5^-^/KRT17^+^ cells, in contrast KRT5^low^/KRT17^+^ co-localised with airway cells. Together, these findings suggest two distinct epithelial origins contributing to fibrotic niches (Figure3K).

**Figure 3.**
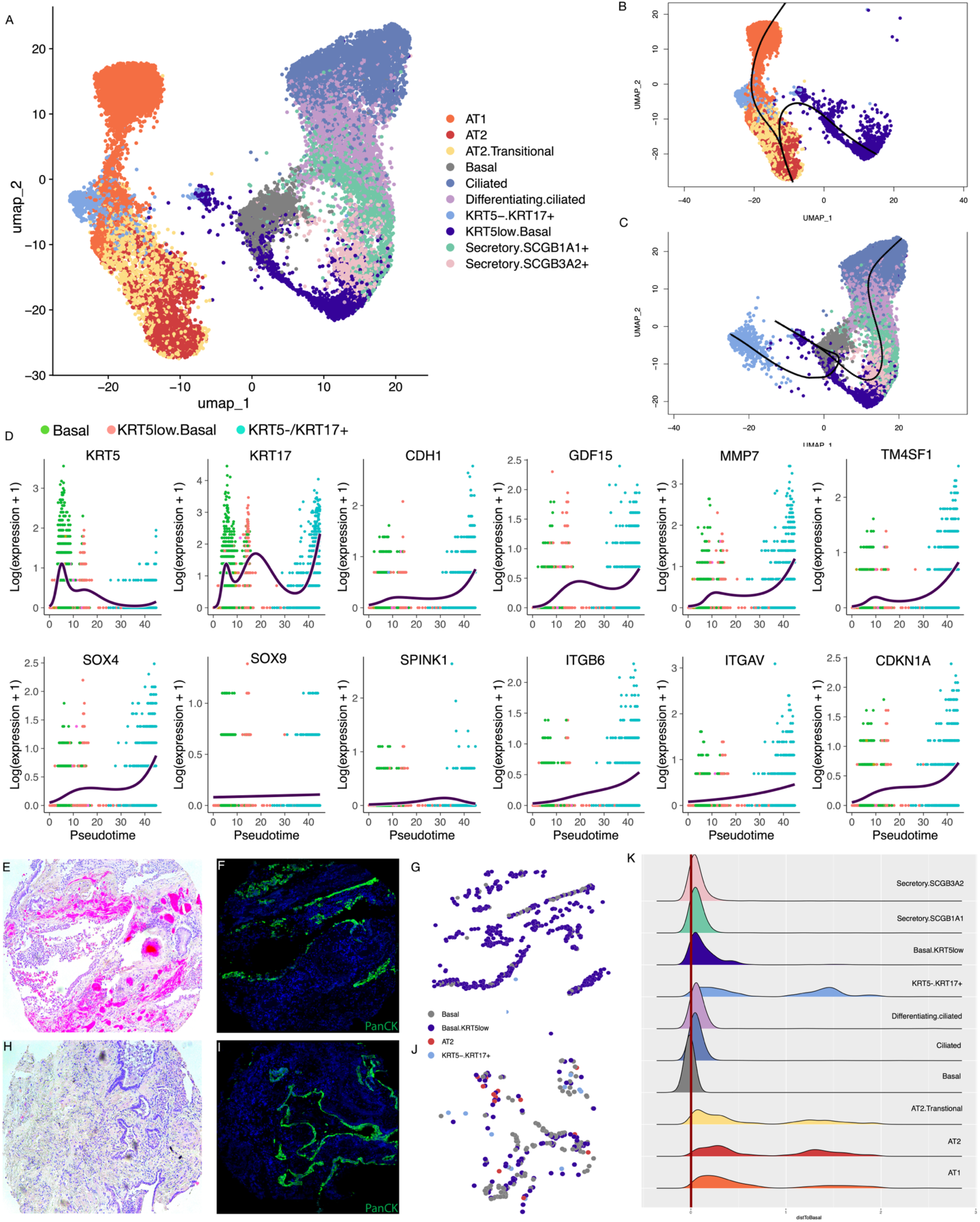
Airway basal cell–derived KRT5^low^/KRT17⁺ transitional state marks progression toward an aberrant basaloid transcriptional signature. **A** UMAP embedding of epithelial subclusters from control and IPF lungs. **B-C** Mapping of KRT5low basal and KRT5⁻/KRT17⁺ aberrant basaloid cells along Slingshot-inferred epithelial lineage trajectories. **B** Alveolar-derived trajectory from AT2 to AT1 and aberrant basaloid states. **C** Airway basal cell–derived trajectory illustrating potential origin of KRT5^low^ and KRT5^-^ cell. **D** Expression dynamics of 12 aberrant basaloid signature genes (*KRT5, KRT17, CDH1, MMP7, SOX4, SOX9, ITGB6, ITGAV, CDKN1A, GDF15, SPINK1, and TM4SF1*) plotted along Slingshot pseudotime across basal, KRT5^low^ basal, and KRT5⁻/KRT17⁺ aberrant basaloid populations. **E,H** Representative hematoxylin and eosin (H&E) images of IPF lung tissues. **F, I** representative CosMx SMI morphology marker (PanCK) images of IPF tissues **G, J** CosMx spatial maps of cell centroids from adjacent tissue sections corresponding to the regions in E and F, highlighting basal cells, KRT5^low^ basal cells, KRT5⁻/KRT17⁺ aberrant basaloid cells, and AT2 cells. I Ridge plots showing the minimal distance of epithelial subtypes to basal cells, calculated from CosMx spatial coordinates. A red vertical line at 0 marks the basal cell location. keratin 5 (*KRT5*), keratin 17 (*KRT17*), cadherin 1 (*CDH1*), matrix metalloproteinase 7 (*MMP7*), SRY-box transcription factors 4 and 9 (*SOX4, SOX9*), integrin subunit β6 (*ITGB6*), integrin subunit αV (*ITGAV*), cyclin-dependent kinase inhibitor 1A (*CDKN1A*), growth differentiation factor 15 (*GDF15*), serine peptidase inhibitor Kazal type 1 (*SPINK1*), and transmembrane 4 L six family member 1 (*TM4SF1*).

### Unique epithelial niches orchestrate aberrant cell–cell communication in fibrotic lung

To investigate how the three distinct epithelial populations contribute to the aberrant micronenvironment, we applied CellChat Ligand-Receptor (LR) analysis to infer cell-cell communication within each niche. Niche analysis revealed three unique epithelial associated niches-N2, an airway niche with cystic airspaces lined by basal cells (“Basal predominant”); N7, an SPP1 macrophage niche enriched for KRT5^low^ basal cells (“Basal.KRT5low predominant”); and N3, the aberrant basaloid niche defined by high level of KRT5^-^/KRT17^+^ epithelial cells (Figure2A, 4A-C). L-R analysis revealed distinct interaction patterns: “Basal predominant” niches showed active crosstalk among airway epithelial cells, macrophages and vascular endothelial cells (Figure4D); “Basal.KRT5low predominant” niches demonstrated strong interactions between KRT5low basal cells and SPP1+ macrophages (Figure 4E); and aberrant basaloid niches displayed marked interactions with activated myofibroblasts (Figure 4F).

**Figure 4.**
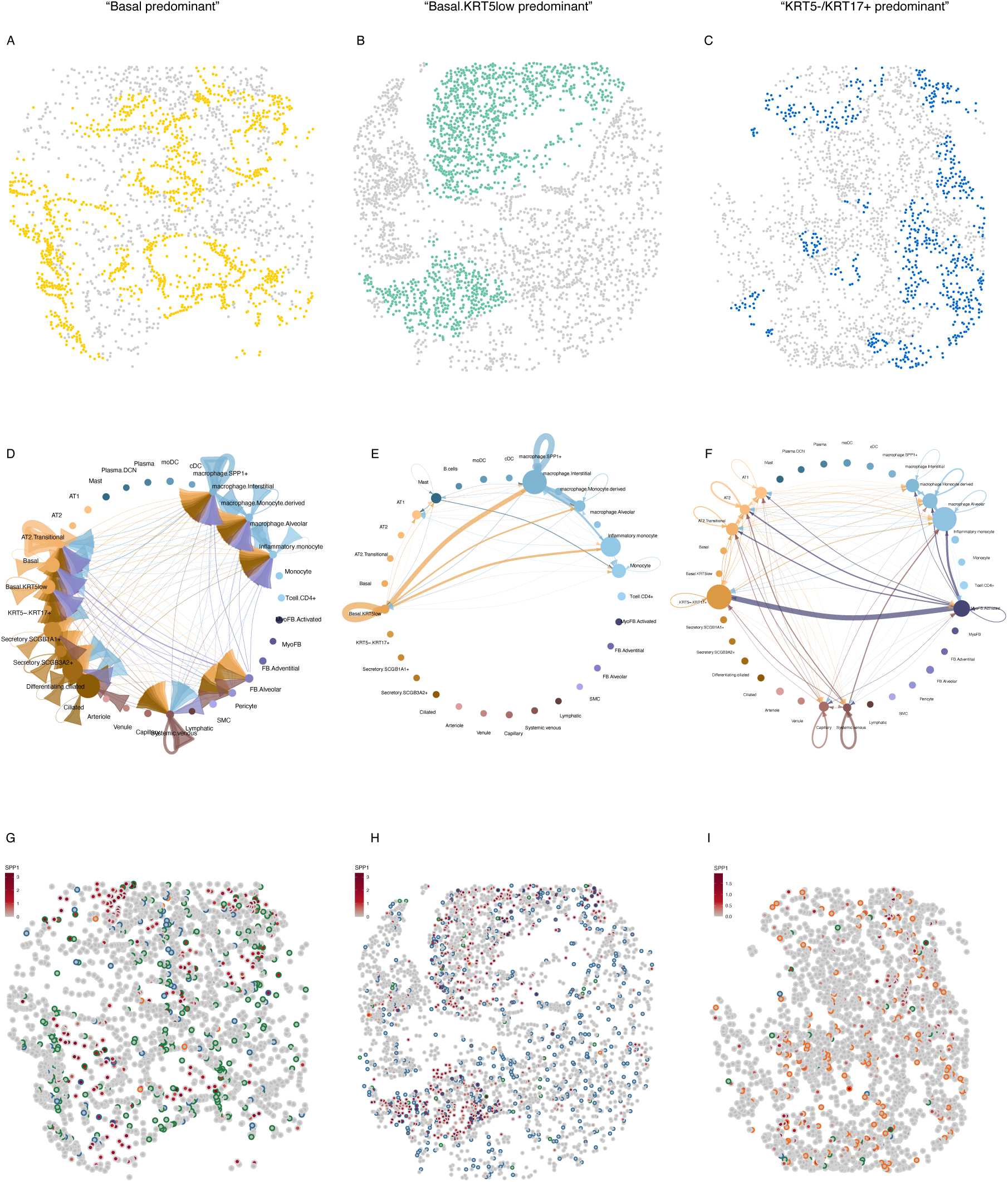
Unique epithelial niches orchestrate aberrant cell–cell communication in fibrotic lung. **A–C** Spatial maps of cell centroids highlighting the location of three epithelial niches: basal-predominant (N2, yellow), KRT5^low^ basal–predominant (N7, green), and KRT5⁻/KRT17⁺ aberrant basaloid–predominant (N3, blue). **D–F** CellChat analysis of inferred ligand–receptor interactions within basal-predominant (D), KRT5^low^ basal–predominant (E), and KRT5⁻/KRT17⁺–predominant (F) niches. Nodes represent cell types and edges indicate predicted interactions, with edge width proportional to interaction strength. **G–I** Co-visualization of cell type centroids and SPP1 expression in basal-predominant (G), KRT5⁻/KRT17⁺–predominant (H), and KRT5^low^ basal–predominant (I) niches. Cell centroid borders are coloured by annotated cell type, and centroid filling is scaled to SPP1 expression levels.

In situ projection of cell centroids onto SPP1 expression maps showed robust overlap of KRT5^low^ basal cell with regions of high SPP1 expression (Figure 4H), compared with basal cell or aberrant basaloid cells (Figure4G, I). Distance measurement confirmed these findings, with KRT5^low^ basal cells showing a sharp peak near zero to SPP1+ macrophages (Figure E4A).

For cross-platform comparison, the KRT5 expression range corresponding to the KRT5^low^ basal population was derived from pseudotime analysis in CosMx data and applied to Hyperion IMC (Figure E4B-C, grey-shaded area), allowing stratification of all KRT17^+^ cells into Basal, KRT5^low^ basal and aberrant basaloid cell populations. Distance analysis revealed a greater proportion of KRT5^low^ epithelial cells localised closer to SPP1^+^ macrophages (Figure E4D), supporting preferential interactions between disease-associated epithelial states and SPP1^+^ macrophages.

To further resolve the potential underlying mechanisms, we specifically examined SPP1-mediated LR pairs across niches. In the basal-predominant (N2) niche, SPP1–(ITGAV+ITGB6) signalling was uniquely restricted to SPP1⁺ macrophages and KRT5^low^ basal cells (Figure E5A). This interaction was further amplified in the basal.KRT5^low^ (N7) predominant niche (Figure E5B), where SPP1⁺ macrophages emerged as the dominant source of SPP1–integrin signaling to KRT5^low^ basal cells. In contrast, the aberrant basaloid niche (N3) displayed a markedly different communication landscape, characterised by broad SPP1-integrin LR pairs, originating from diverse cell types and converging on KRT5^-^/KRT17^+^ cells (Figure E5C). In addition, the immune–stromal niche (N4), composed of lymphocytes, vascular endothelial cells, and alveolar fibroblasts, was characterised by strong ECM–receptor and chemokine signalling (Figure E6).

### Basal cell detachment is associated with loss of KRT5 expression highlighting a novel origin of fibrotic epithelial cells

Previous transcriptomic studies have not identified a population of KRT5^low^ cells derived from basal cells, however work by Vanna and colleagues(22) revealed that epithelial cell detachment was closely associated with transcript based niches that associate with KRT5-/KRT17+ cells. Therefore, to understand whether epithelial detachment could lead to loss of KRT5, patterns of KRT5 and KRT17 expression were assessed in overcrowded monolayers of iHBECs and primary basal cells and in the cells which detached from these monolayers. Detached iHBECs showed a dramatic reduction in KRT5 mRNA but retained high levels of *KRT17* mRNA (Figure 5A). Flow imaging cytometry of individual iHBECs confirmed that cells that had been retained in the monolayer had consistently high levels of KRT5 protein which was reduced or absent in iHBECs that had detached from the monolayer (Figure 5B). Similarly, flow cytometric analysis showed expression of KRT5 and KRT17 were both high in iHBEC (Figure 5C) and primary basal cell monolayers (Figure 5D) but there was a distinct population of KRT5^-^ and KRT5^low^ cells in both iHBECs and primary basal cells following detachment (Figure 5C and D). Finally to understand if mechanical forces might contribute to the development of the fibrotic epithelial cell phenotype, primary small airway epithial cells were subjected to cyclical mechanical stretch for upt to 24hrs and showed a progressive loss of KRT5 expression while KRT17 remained intact (Figure 5E).

**Figure 5.**
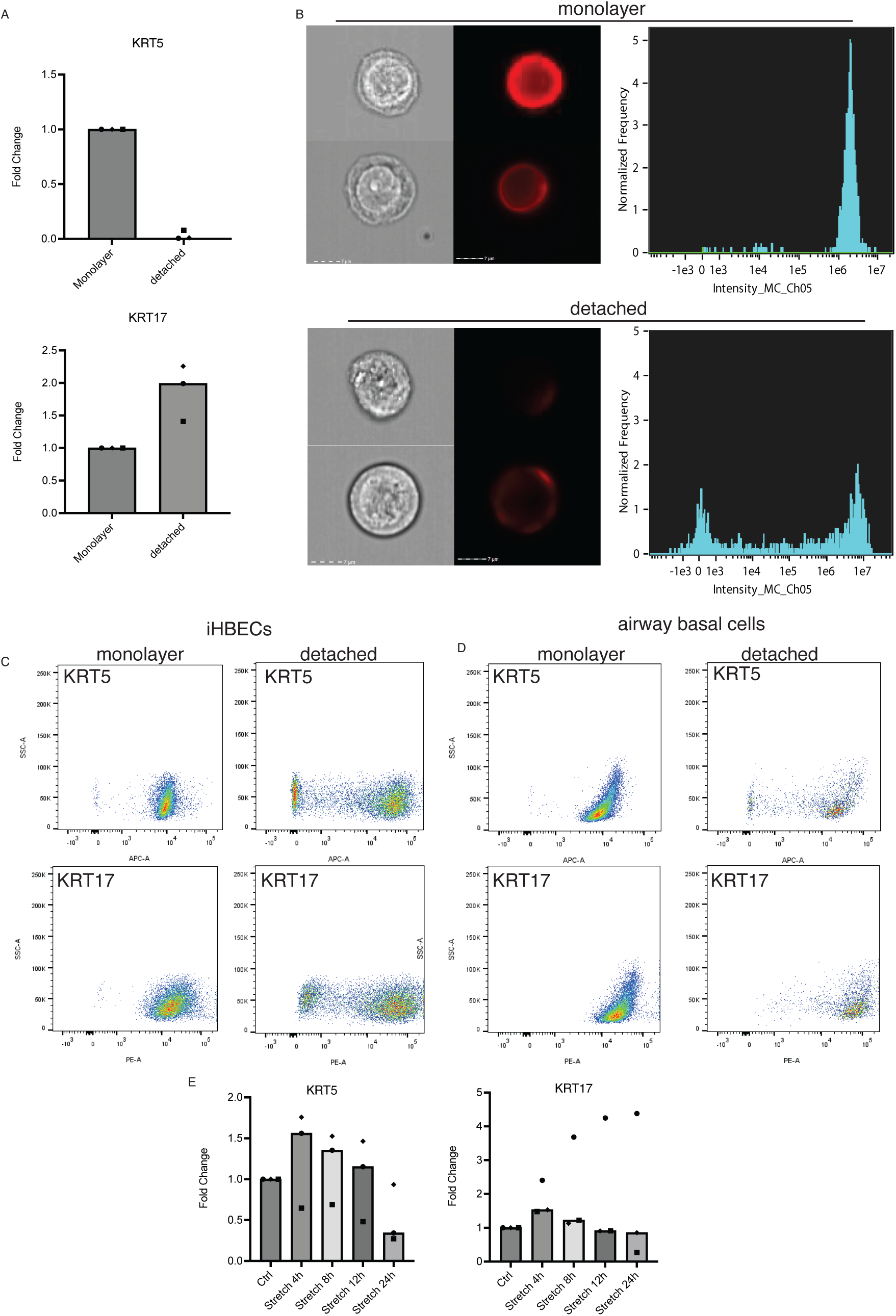
Basal cell detachment is associated with loss of KRT5 expression. **A** ihBECs mRNA Expression of KRT5 and KRT17 (n=3) of monolayer cells and detached cells (collected from supernatant) **B** ihBECs ImageStream MK II Imaging Flow Cytometry of monolayer cells and detached cells (representative data n=2) stained with intracellular KRT5 antibody (2ug/mL). **C** Flow Cytometry using BD FACSCanto of ihBECs monolayer cells and detached cells stained for KRT5 (APC-A) and KRT17 (PE-A) at 2ug/mL (representative data n=3). **D** Flow Cytometry using BD FACSCanto of primary basal monolayer cells and detached cells stained for KRT5 (APC-A) and KRT17 (PE-A) at 2ug/mL (representative data n=2). **E-F** *KRT5* and *KRT17* mRNA expression of small airway epithelial cells (SAECs) subject to cyclic stretch over 24 hours (n=3).

## Discussion

This study applied bi-level spatial multi-omics to resolve the complex cellular architecture of IPF lung tissue, delineate the spatial organization and interactions of key IPF-associated cell populations, and define distinct fibrotic niches. Our findings confirm and extend single-cell(4, 5, 7, 8) and recent spatial transcriptomic studies(22–24) that identified IPF-enriched populations such as KRT5⁻/KRT17⁺ aberrant basaloid cells, activated myofibroblasts, and SPP1⁺ macrophages and described dysregulated niches characterized by alveolar loss, macrophage reprogramming, and fibroblast activation. Importantly, here we reveal an intermediate KRT5^low^/KRT17⁺ basal population enriched within fibrotic niches, demonstrating experimentally that epithelial detachment and mechanical stretch result in KRT5 loss, with these cells localized within SPP1⁺ macrophage-enriched fibrotic regions.

Consistent with prior murine(7, 25) and human (6, 26) single-cell studies, our trajectory analysis suggests that KRT5^-^/KRT17^+^ aberrant basaloid cells could arise from AT2 cells and form a fibrotic niche in association with activated myofibroblasts. Intriguingly, our analysis also identified an intermediate transcriptional profile of the KRT5^low^/KRT17⁺ population between KRT5^+^ basal and KRT5⁻/KRT17⁺ aberrant basaloid cells. Distance analysis reveals a close but not overlapping spatial location of these KRT5^low^ transitional cells to basal cells, suggesting their origin. In murine models, lineage tracing has demonstrated that KRT5⁺ airway basal cells can migrate into the alveolar region following injury(27, 28), whereas in human lung they have been implicated in IPF pathogenesis through de-differentiation into KRT17⁺ epithelium(29), and aberrant expansion into the distal lung under fibrotic ECM remodeling(30). Our data further support that KRT5⁺ basal cells comprise a substantial fraction of epithelial cells within the fibrotic niche although the role they play in fibrogenesis is not clear. Airway basal cells have been previously identified as self-renewing progenitor cells capable of differentiating into various lung epithelial cells(31–34) following injury, and this observation suggests that KRT5⁺ basal cells may also serve as a source for the KRT17⁺ fibrotic epithelium, supporting the hypothesis that small airways could play a central role in IPF pathogenesis(35). Whether this reflects a primary defect following airway injury, or a secondary response to alveolar injury, collapse and fibrotic contraction leading to traction associated change of the small airways remains unclear. A recent study has also shown that epithelial detachment overlaps with niches enriched for KRT5^-^/KRT17^+^ aberrant basaloid cells(22). In line with this, our *in vitro* data confirm that detachment, and mechanical forces, can trigger progressive loss of KRT5 expression across diverse airway epithelial cells. Taken together, these findings suggest that in the disease state, airway basal cells may transition through a KRT5^low^/KRT17^+^ intermediate population, acquiring aberrant basaloid transcriptional profile, and contribute to the formation of fibrotic niches.

Our identification of three distinct epithelial niches raises the intruiging possibility that loss of both AT1 and AT2 cells may be the initial defect in fibrogenesis. As AT2 cells fail to transdifferentiate appropriately, aberrant basaloid cells accumulate and form a primary fibrotic niche in association with activated myofibroblasts. Progressive matrix deposition and myofibroblast expension within this niche likely alters mechanical force generation around the small airways, driving the emergence of two “secondary fibrotic niches”. The first is a basal cell predominant niche characterised by dilated and expanded airways(30), and the second is a KRT5^low^/KRT17^+^ niche, marked by detached and stretched epithelium that progressively loses KRT5 in an attempt to compensate for the loss of alveolar epithelial cells. With disease progression, these secondary fibrotic niches expand, as reflected by increasing numbers of both KRT5^+^ basal cells and KRT5^-^/KRT17^+^ cells in more severe IPF^27^. Notably, niche analysis indicates that KRT5^low^/KRT17^+^ form a distinct niche in close association with SPP1^+^ macrophages raising the interesting possibility that interaction of SPP1^+^ macrophages, either directly or indirectly, with airway basal cells through the SPP1-αvβ6 integrin axis can promote pulmonary fibrosis. However future functional studies are warranted to confirm this potential interaction.

In addition our analysis also highlights the emergence of distinct immune–stromal niches in IPF. We observed a subset of lymphocytes preferentially localise within decorin-rich stromal regions, where they interact strongly with vascular endothelial cells and alveolar fibroblasts. Ligand–receptor inference revealed that alveolar fibroblasts signal to plasma cells through CXCL12–ACKR3, and collagen–CD44 pairs, suggesting a matrix-dependent mechanism of plasma cell recruitment and retention. A recent study by Yang et al.(36) demonstrated that CXCL12-expressing fibroblasts promote plasma cell accumulation within tertiary lymphoid structures in IPF, consistent with our observations. These observations highlight dynamic immune–stromal crosstalk in IPF, the functional consequences of which remain to be fully understood.

Our study represent one of the largest spatial multi-omics comparisons of IPF and control lungs, with transcriptomic findings validated at the protein level by Hyperion IMC. We also provide proof-of-principle that airway basal cells can lose KRT5 expression under conditions relevant to fibrogenesis. However, our study has some limitations. The nature of spatial transcriptomics limits the number of samples available for analysis and impacts the generalisability of our findings, although the replication of previously described niches is reassuring. Similarly there was incomplete coverage of transcripts for analysis, with CTHRC1 being absent from the CosMx SMI, however, replication with Hyperion IMC and the use of alternative markers increase confidence in our findings.

In conclusion, this study identified three distinct fibrotic epithelial niches: a primary fibrotic niche characterised by aberrant basaloid (KRT5^-^/KRT17^+^) cells and activated myofibroblasts; and two secondary fibrotic epithelial niches characterised by airway basal cells or basal cell derived KRT5^low^/KRT17^+^ population in close association with SPP1^+^ macrophages. The enrichment of KRT5⁺ airway basal cells and abnormal KRT5^low^/KRT17⁺ basal cells within distinct niches supports the hypothesis that airway basal cells serve as a major source of fibrotic epithelium. Together with immune–stromal niches that may contribute to fibroblast activation, these findings provide a refined cellular framework for understanding the evolution of progressive fibrotic lung disease.

## Sources of supports

This study was funded by a Medical Research Council Programme Grant (MR/V00235X/1) to R.G.J and MRC Multi-modal grant (MR/W031469/1) awarded to R.G.J, A.E.J, I.D.S., C.D., M.P. and J.R.. RGJ was funded by an NIHR Research Professorship (RP-2017-08-ST2-014). B.L. is a research fellow funded by Action for pulmonary fibrosis. I.D.S. is an advanced research fellow funded by Rayne Foundation. Infrastructure support for this research was provided by the NIHR Imperial Biomedical Research Centre (BRC). The authors acknowledge the Nottingham Biomedical Research Centre (BRC) and Clinical Research Facility (CRF) Respiratory Biobank, Royal Brompton Hospital for their collection of tissue and facilitation of sample transfer. The authors also thank Prof. Paul Matthews, Vicky Chau, and Dorcas Cheung for their technical support in the application of Hyperion imaging mass cytometry.

## Author contributions

B.L., K.B., I.D.S., J.M., M.C.Z., I.U., and E.L.-J. contributed to tissue sample collection, in vitro experiments, and data analysis. S.R.J., A.L.T., and R.L.C. provided critical scientific input and supplied human lung tissues. B.L., N.L., and N.M. conducted CosMx sample preparation, processing, and data acquisition. M.P., J.R., R.H., L.V.W., A.K.R., E.A.R., P.M.G., P.L.M., C.D., and A.U.W. critically reviewed and revised the manuscript. B.L., A.E.J., and R.G.J. conceptualized the study and drafted the manuscript, with K.B. contributing to manuscript preparation. A.E.J. and R.G.J. supervised the project, coordinated collaborations, and finalised the manuscript. All authors reviewed and approved the final version of the manuscript.

