## supplemental materials and methods for "Spatial multiomic profiling reveals distinct fibrotic epithelial niches in idiopathic pulmonary fibrosis"

Bin Liu *et al.*

**This PDF file includes:**

Materials and Methods

### Materials and Methods:

#### Human tissue collection and tissue microarray construction

Human lung tissue blocks from IPF patients were sourced from the Nottingham NIHR Biomedical Research Centre under ethical approval (REC 08/H0407/1). Non-IPF tissues were freshly collected from Clinical Research Facility (CRF) Respiratory Biobank, Royal Brompton Hospital under the ethical approval (NRES 20/SC/0142). Lung tissue was formalin-fixed and paraffin-embedded prior to their use in constructing a tissue microarray (TMA) block. For the TMA construction, 1 mm cores were sampled from IPF (4 cores) and control (2 cores) tissues.

#### Slides preparation and Data acquisition:

Formalin-fixed, paraffin-embedded (FFPE) tissue blocks were sectioned at a thickness of 5 µm. Sections were mounted on VWR® SuperFrost® Plus slides in the CosMx scanning area (20mm by 15mm) and were stored in a sealed box with desiccants at 4°C to maintain dryness and prevent degradation.

For the CosMx Molecular Imager, slides were baked at 60°C to promote optimal tissue adherence. Slides were then deparaffinised followed by target retrieval at 100°C, tissue permeabilization, and digestion with 3ug/ml digestion buffer for 30 minutes. Post-fixation in 10% Neutral Buffered Formalin (NBF) was subsequently performed, followed by N-Hydroxysuccinimide (NHS)-Acetate incubation prior to labelling. Overnight hybridization with the Human Universal Cell Characterization Panel (1000-plex RNA, Bruker, USA) was carried out at 37°C. Following stringent washing, nuclear staining with DAPI and cell morphology maker staining with pan-cytokeratin (PanCK)/CD45, together with segmentation markers CD298/B2M, were performed using Human RNA Universal Cell Segmentation kit (Bruker, USA), for cell segmentation and downstream analysis. Configuration C was used as pre-bleaching profile.

For Hyperion imaging mass cytometry, carrier-free antibody metal conjugation was performed using the MaxPar antibody labeling kit following the manufacture protocol (Standard BioTools, USA). All labelled antibodies were eluted in antibody stabilizer (Candor Bioscience, Germany) supplemented with 0.05% sodium azide. FFPE sections were baked for 2 hours at 60°C. Dewaxing was performed using fresh Histo-Clear (A2-0101, geneflow) for 20 minutes, followed by a hydration sequence through descending ethanol concentrations. The antigen retrieval was performed in Citrate Buffer (ab93678, abcam) at 96°C for 20-minutes. Following phosphate buffered saline (PBS) wash, tissue sections were blocked with 3% Bovine Serum Albumin (BSA) in PBS for 60 minutes at room temperature. An antibody cocktail, prepared according to specific concentrations (Table E1) was applied and sections were incubated overnight at 4°C. Following washing tissue sections were incubated with Intercalator-Ir in PBS for 30 minutes at room temperature.

**Data processing and single cell Data Analysis:**

**CosMx SMI data processing and analysis**

Machine learning-driven single cell segmentation was performed using AtoMx Spatial Informatics Platform (SIP)(E1, E2). For data quality control (QC), cells with negative probe greater than 0.5 and or features less than 20 were filtered out. In total, 180,067 cells were profiled, and raw counts of all cells were exported as a Seurat object from AtoMx SIP for downstream analysis. Total counts normalization was applied. Following normalization, principal component analysis (PCA) and then Uniform Manifold Approximation and Projection (UMAP) and t-distributed Stochastic Neighbor Embedding (tSNE) were performed for dimension reduction and visualisation using Seurat v5(E3).

**Cell type annotation**

We applied a hierarchical strategy for cell type annotation, starting with Leiden clustering and aggregating clusters into the main cell lineages based on Human Lung Cell Atlas (HLCA) v2(E4) level1 markers which are included in the CosMx 1000-plex gene panel: Immune markers: (CD53, PTPRC, COTL1, CXCR4, FCER1G, SRGN, CD52); Epithelial markers: (KRT7, PIGR, KRT8, KRT19, TSPAN8, CSTD2, CXCL17); Endothelial markers: (PECAM1, VWF, RAMP2, IGFBP7, CLEC14A); and Stromal markers: (TPM2, DCN, MGP, CALD1, LUM, TAGLN, COL1A1). These four lineage subsets were then used for downstream cell subtype annotation using the Insitutype algorithm(E5), applying semi-supervised cell typing. Briefly, a cell profile matrix was firstly derived from public available single cell RNAseq data (GSE227136) using SpatialDecon(E6) and further used to identify “anchor cells" with confident reference cell type assignments. The mean profiles of these anchor cells were then used to refine our reference matrix, ensuring a more accurate representation of cell types in the CosMx context. To enrich the analysis, cells were organised into cohorts based on immunofluorescence values of the morphology markers (PanCK and CD45) using fastCohorting function within Insitutype package.

**Inference of inter-niche and inter-cell communication using CellChat V2**

For inference of cell communication, we first constructed a Niche assay from CosMx spatial transcriptomics data by aggregating local cell-type composition and grouping similar neighbourhoods into recurrent niches. Clustering of neighbourhoods revealed seven distinct niches (N1–N7). N1: MyoFB niche, N2: Airway niche,N3: KRT5-/KRT17+ niche,N4: Immune niche,N5: Vascular niche,N6: Alveolar niche, N7: SPP1+ macrophage niche. CellChat V2(E7) was used to infer spatially proximal inter-niche communication and intra-niche intercellular communication. CellChat objects were created separately from the Seurat object. CellChat DB V2 was used as the ligand-receptor (L-R) interaction database. A total of 358 L-R pairs were extracted from the NanoString 1000-plex RNA probes panel and used for further analysis. Both control and IPF CellChat objects were then merged and compared for total interaction number and strength using the compareInteractions() function. Interactions between cell populations were visualized in circle plots using the netVisual_diffInteraction() function. The cell-cell communication network of key fibrotic cell types was visualized separately. To further investigate pathway-level dysregulations, significant LR pairs contributing to niche- or cell- specific communication were aggregated into annotated pathway within the CellChat package and visualised using rankNET() function. Individual LR pairs that contribute to the communication were further visualised by bubbleplot for interaction strength and significance.

**Hyperion IMC data processing and analysis**

**Image preprocessing, segmentation and feature extraction**

Preprocessing, segmentation, and feature extraction of multi-channel Hyperion IMC images were achieved using the IMC Segmentation Pipeline, as previously described(E8). Briefly, raw .mcd files were first converted into .ome.tiff files using the imctools Python package. A random forest-based image pixel classification was performed using ilastik. Pixels classified as "cytoplasm", "nucleus", and "background" were input for probability assessments and then thresholded using the CellProfiler pipeline to generate segmented cell masks. Features including the mean pixel intensity per cell and channel, cell morphological features, and locations were extracted using a customised CellProfiler pipeline for downstream analysis.

**Single-cell spatial analysis of IMC data**

Image based single cell spatial analysis of the IMC data was performed using imcRtools and cytomapper R package. Briefly, SpatialExperiment object was created using imcRtools while multi-channel images and cell masks were loaded using cytomapper. Unsupervised cell clustering was performed using Rphenograph. Graph based clustering was used to generate shared nearest neighbour (SNN) graphs. Cell type classification was carried out based on pre-defined markers, with fibrotic epithelial cells as KRT17+/KRT5-, fibrotic fibroblasts as CTHRC1+, fibrotic macrophages as SPP1+, and fibrotic B cells as CD20+/DCN+. To identify fibrotic associated cellular niches, cellular neighbourhood analysis was performed to compute the 20-nearest neighbours and kmeans=5 for clustering cells into 5 cellular niches.

**Cell culture**

Immortalised human bronchial epithelial cells (iHBECs) were mained in Keratinocyte-Serum-Free-Medium (Gibco) containing L- glutamine supplemented with 25µg/mL Bovine Pituitary Extract (Life Technologies), 0.2ng/mL human recombinant Epithelial Growth Factor (Life Technologies), 250ng/mL Puromycin (Merck) and 25µg/mL Geneticin Selective Antibiotic (G418 Sulfate) (Life Technologies). Primary basal cells were isolated from human lung tissues. Briefly, Fresh tissue sample was cut into 1-2mm diameter size biopsy and plated 20-30 per dish into collagen coated petri dishes. The biopsies were left to adhere for 10mins before adding 4ml of airway epithelial cell growth media. Tissues were maintained for 7-10 days until 50% of biopsies released basal cells before removal to reduce fibroblast contamination. Basal cells were used at P3 for experiment. For inducing detachment, cells were seeded at a high density (2 million cells per T175) to cause overcrowding. Once confluency was reached, cells were maintained in cluture for a further week, detached cells were collected and dead cells were removed using Miltenyi Dead Cell Removal kit. Human small airway epithelial cells(SAECs) obtained from Lonza were cultured in Clonetics™SAGM™BulletKit™ (CC-3118) following the manufacturers protocol and subject to 15% elongation at 0.3Hz of cyclical stretch (Flexcell FX5K Tension, Dunn Labortechnik) over 24 hours period.

**RNA Extraction and Real-time qPCR**

RNA extraction was performed using Maxwell® RSCExtraction System (AS1390, Promega, USA) following manufacturer protocol or with the NucleoSpin™ RNA Mini Kit (Macherey-Nagel, #12373368) following the manufacturers protocol. 100ng of RNA isolated from monolayer cells or detached cells from ihBECs was used in the cDNA synthesis. *KRT5* (Primer Forward: GCTGCCTACATGAACAAGGTGG and Primer Reverse: ATGGAGAGGACCACTGAGGTGT) and *KRT17* (Primer Forward: ATCCTGCTGGATGTGAAGACGC and Primer Reverse: TCCACAATGGTACGCACCTGAC) genes expression was measured with real-time qPCR using SYBR GREEN Fast Plus system. Geometric mean of housekeeping genes (*HPRT1* and *B2M*) were used for normalisation.

**Imaging Flow Cytometry**

Imaging flow cytometry of KRT5 in ihBECs was performed using the ImageStream MK II system. Cells were permeabilised and fixed using eBioscience™ Foxp3 / Transcription Factor Staining Buffer Set (Thermo Fisher Scientific) following the supplier's protocol. Following fixation, cells were washed and blocked for 30mins at 4° in the dark before stained for KRT5 (2ug/mL) antibody for 30mins at 4° in the dark. Cell were then resuspended in 50ul FACs buffer (0.5% BSA and 0.1% Sodium azide in PBS) after straining through a 40um cell strainer to achieve a single cell suspension. Gating and downstream analysis was performed using IDEAS software. For gating, aspect ratio (defined by the ratio between the longest and shortest line that can be drawn through the shape) versus the area of the brightfield mask were used. Cells with medium area and high aspect ratio were kept to remove cell clusters and speed beads/debris as shown in Figure E1.

**Flow Cytometry**

Flow cytometry assessment of KRT5 or KRT17 expression in ihBECs and primary Basal cells was performed using the BD FACSCanto system and analysed using FlowJo. The cells were permeabilised and fixed following the same method as detailed in Flow Imaging Cytometry section. Cells were co-stained for KRT5 and KRT17 (2ug/mL). Final volume was suspended in 100ul FACs buffer and 200ul of PBS was added to each sample before analysed by the flow cytometer. Gating and analysis was performed using FlowJo. The unstained sample was the negative control for the gating and same parameters were then applied to the co-stained detached and monolayer samples for KRT5 and KRT17 levels comparability as shown in Figure E1.
