## supplemental figure1-6 and table1 for "Spatial multiomic profiling reveals distinct fibrotic epithelial niches in idiopathic pulmonary fibrosis"

Bin Liu *et al.*

**This PDF file includes:**

Figure E1 to E6
Tables E1

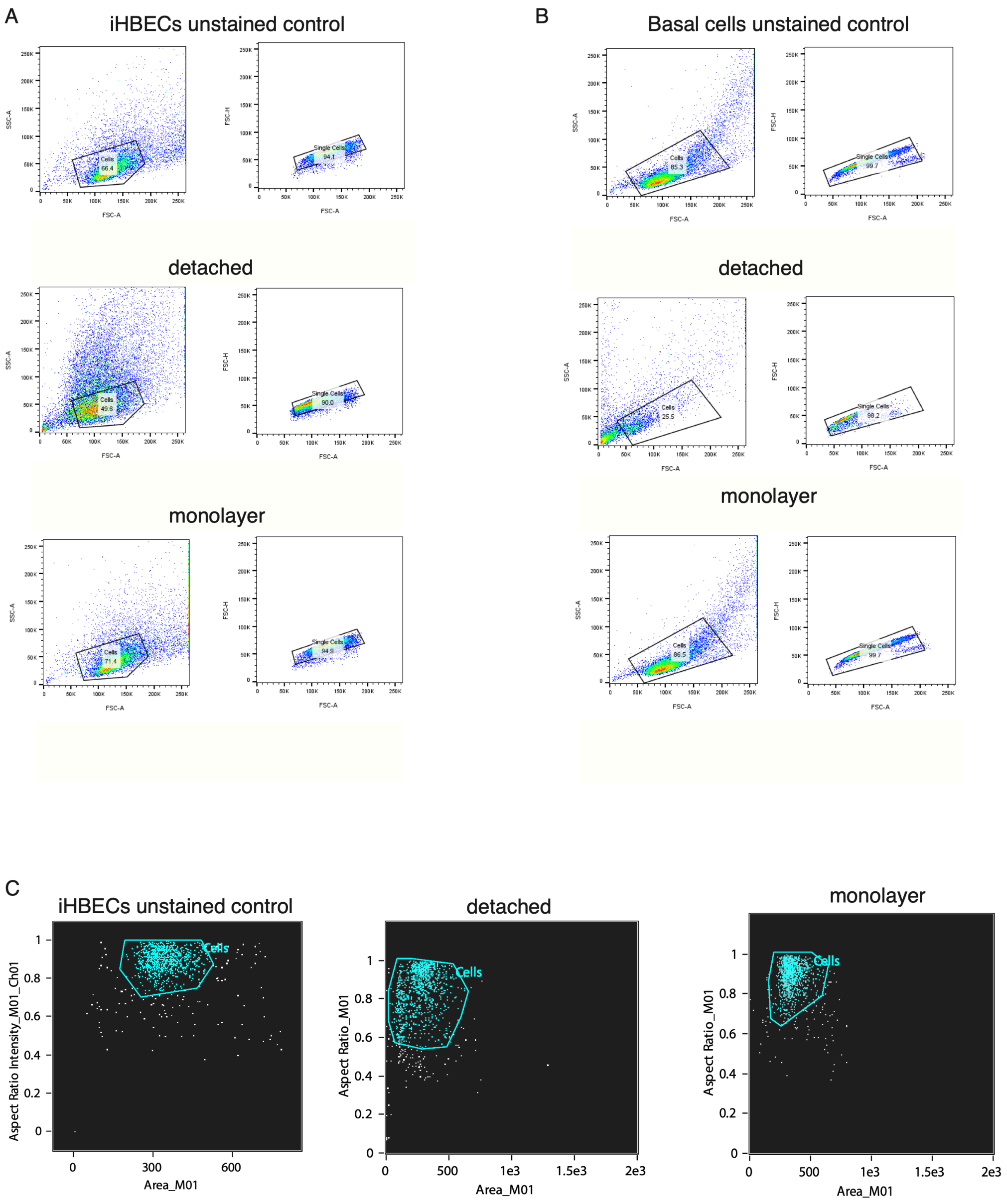


**Figure E1 Gating strategy for Flow cytometry and Imaging Flow cytometry** Scatter plots of flow cytometry data illustrating gating strategies for (**A**) iHBECs and (**B**) primary basal cells. For each, the top row shows unstained cells, the middle row detached cells, and the bottom row monolayer cells. Side-scatter area (SSC-A) versus forward-scatter area (FSC-A) was used to select the main cell population, while forward-scatter height (FSC-H) versus FSC-A was used to exclude doublets. (**C**) Scatter plots of Aspect Ratio Intensity versus Area used to visualise the gating strategy for Imaging flow cytometry.

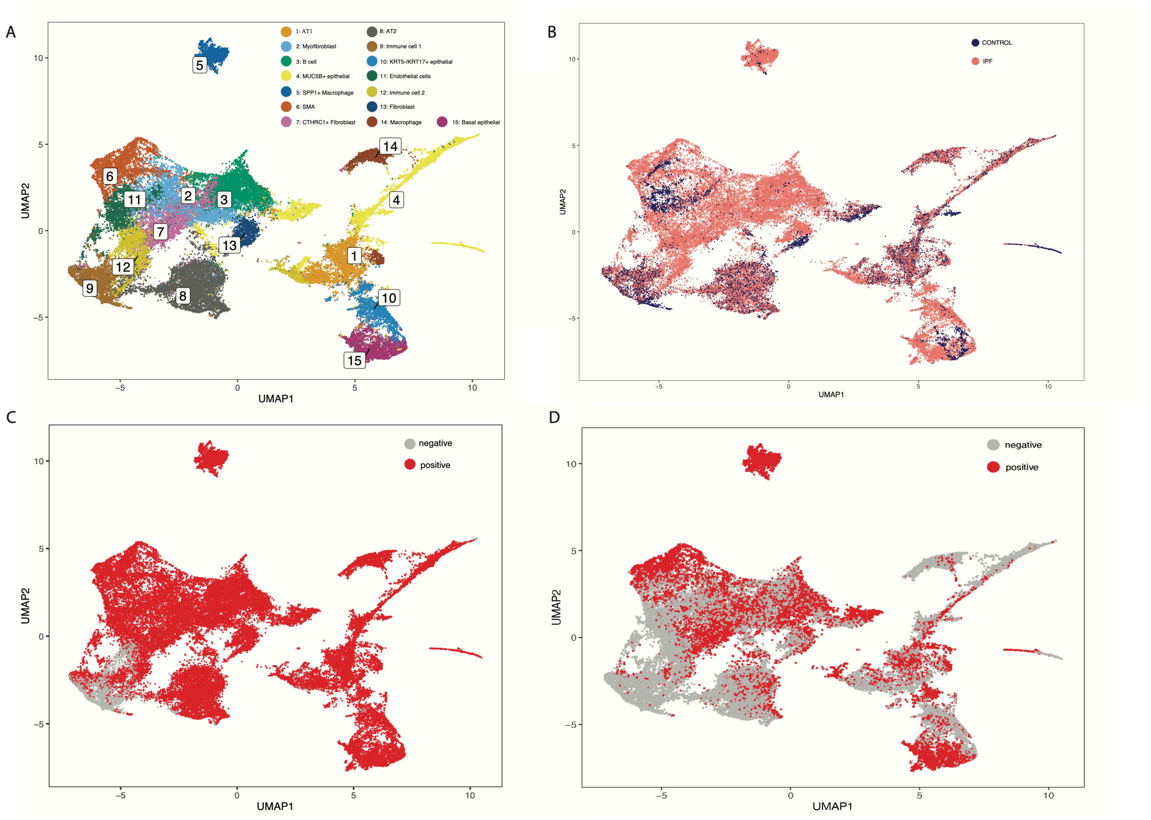


**Figure E2. Cross-platform comparison of cell populations in IPF and control lungs using Hyperion IMC** UMAP embedding visualisation of **A** 15 cell types identified, **B** the same embedding coloured by disease status (control versus IPF), **C** cells expressing at least one of the shared **markers (NDRG1, VIM, EPCAM, SPP1, PECAM1, PTPRC, KRT17, COL4A1, PDGFRB, CD68, KRT5, CDKN1A, ARG1, DCN, MKI67, COL1A1, COL15A1, KRT8)** across CosMx and Hyperion (red, positive; grey, negative), and **D** cells expressing at least one of the shared **fibrotic markers (KRT5, KRT17, SPP1, CTHRC1, DCN)** across the two platforms (red, positive; grey, negative).

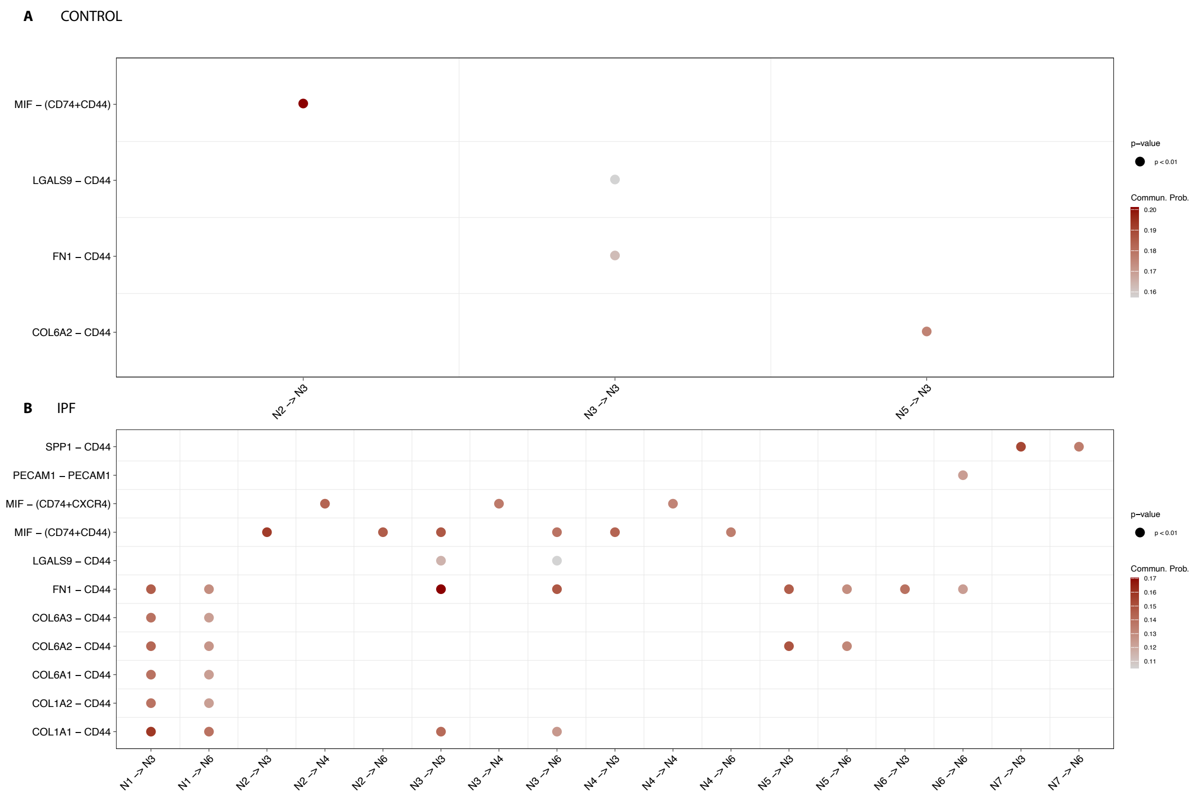


**Figure E3. Niche composition and communication analysis of CosMx SMI data of control and IPF lung** Bubble plots of ligand–receptor pairs across niches N1–N7 in control (top) and IPF (bottom).

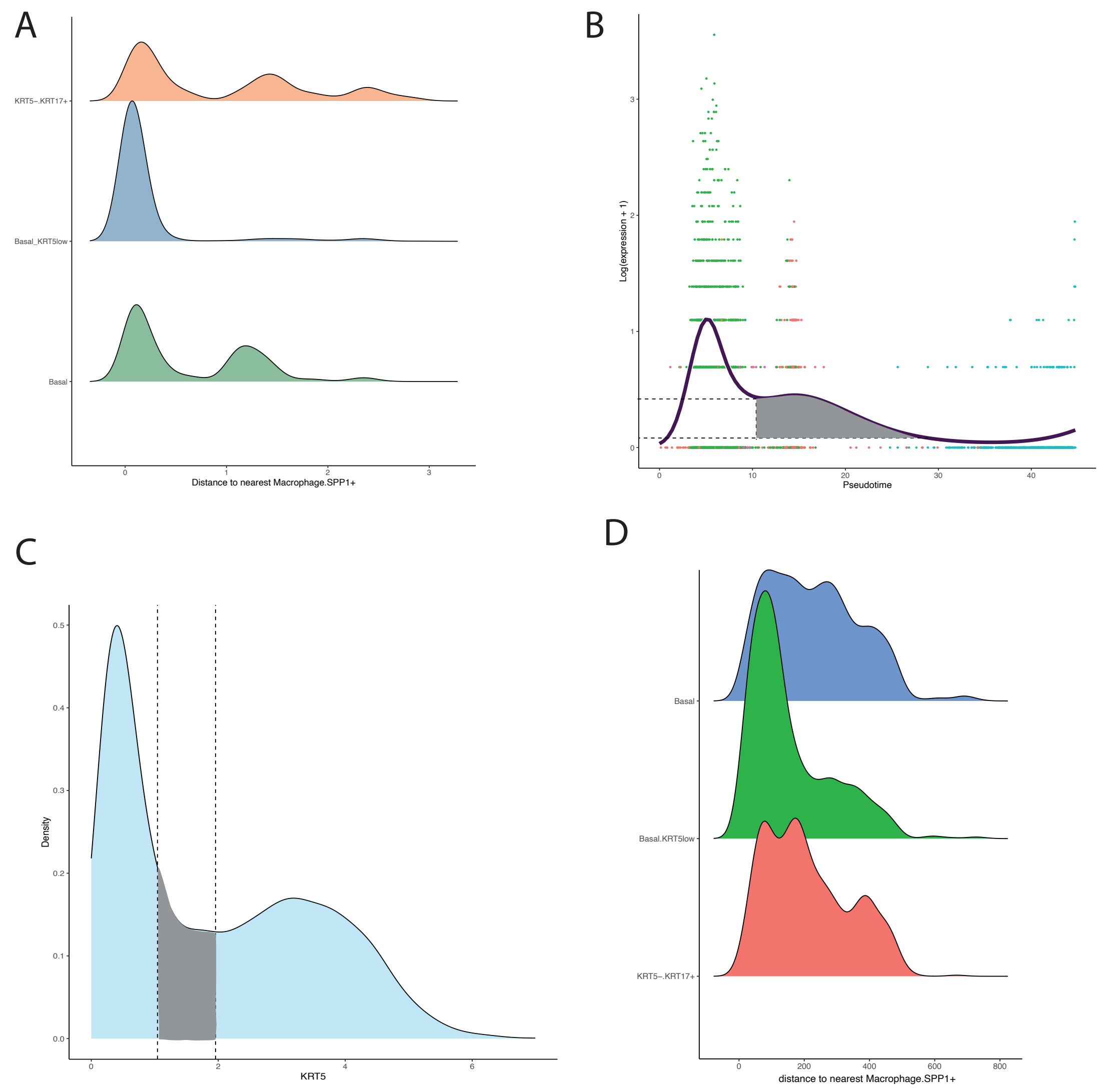


**Figure E4 Spatial proximity of KRT17+ epithelial cells to SPP1$^+$ macrophages**

**A** Ridge plots of minimal distance to SPP1$^+$ macrophages for basal, KRT5$^{low}$ basal, and KRT5$^-$/KRT17$^+$ epithelial cells calculated using CosMx spatial coordinates. **B** KRT5 expression along pseudotime in CosMx, with the grey-shaded region marking the expression range defining KRT5$^{low}$ basal cells. **C** Density distribution of KRT5 expression in Hyperion IMC, stratified using thresholds guided by the CosMx-derived KRT5$^{low}$ range in panel B (grey-shaded). **D** Minimal distance to SPP1$^+$ macrophages for the stratified epithelial populations in Hyperion IMC.

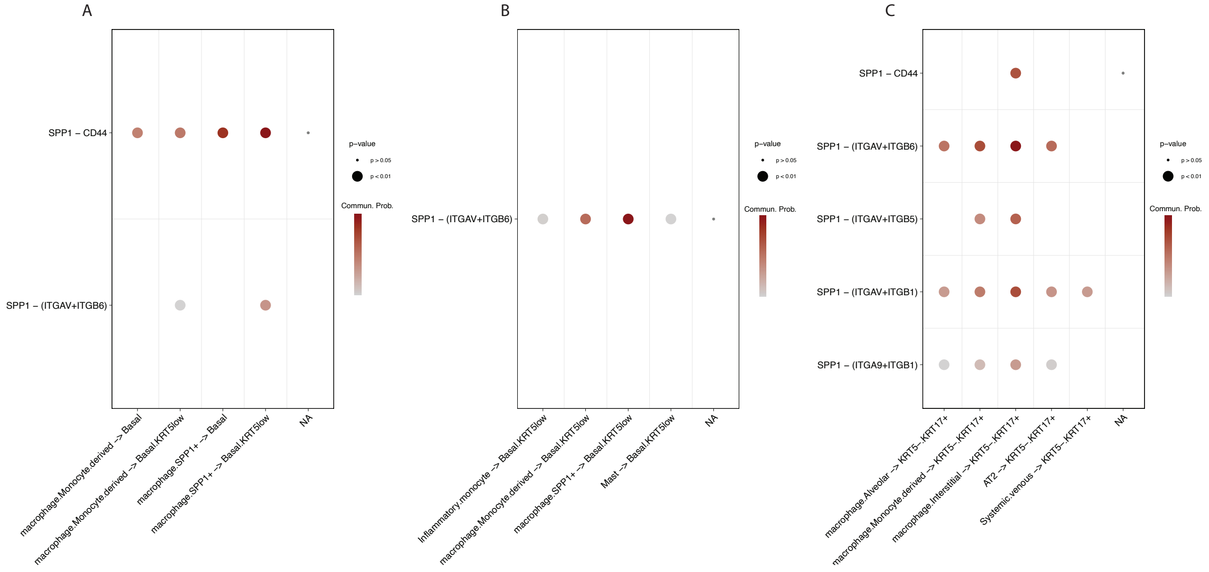


**Figure E5 SPP1 pathway ligand–receptor interactions across epithelial-associated niches.** Bubble plots from CellChat showing SPP1-related ligand–receptor pairs in **A** the basal-predominant niche, **B** the Basal.KRT5$^{low}$-predominant niche, and **C** the aberrant basaloid niche. Bubble color denotes the communication probability, and bubble size reflects the statistical significance p value.

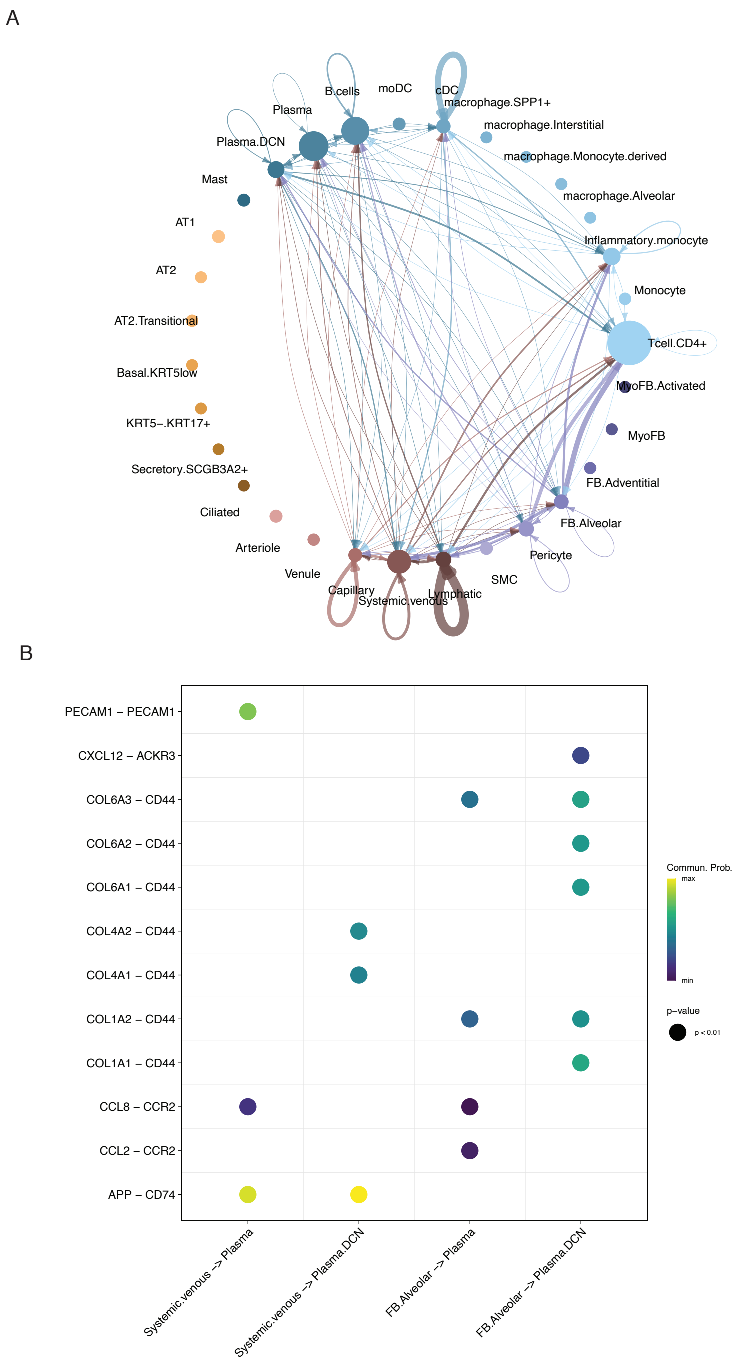


**Figure E6 Cell–cell communication in the immune–stromal niche. A** Circle plot of CellChat-inferred interactions within the immune–stromal niche (N4). Node size reflects the relative abundance of each cell population, and edge thickness denotes the overall strength of predicted interactions between populations. **B** Bubble plots of ligand–receptor pairs from the top enriched pathways among lymphocyte, endothelial, and fibroblast subtypes. Bubble colour indicates communication probability, and bubble size represents statistical significance (p value).

| TARGET | METAL | DILUTION | CLONE |
|---|---|---|---|
| SMA | 141 Pr | 1/1000 | 1A4 |
| NDRG1 | 142 Nd | 1/200 | PA5-80847 |
| Vimentin | 143 Nd | 1/1000 | D21H3 |
| EPCAM | 144 Nd | 1/100 | EPR20532-222 |
| SPP1 | 146Nd | 1/100 | 7C5H12 |
| MUC5B | 147 Sm | 1/100 | ab87376 |
| CTHRC1 | 148 Nd | 1/50 | EPR22851-145 |
| Keratin17 | 149 Sm | 1/200 | EPR1624Y |
| P21 | 150 Nd | 1/100 | WA-1 |
| CD31 | 151 Eu | 1/400 | EPR3094 |
| CD45 | 152 Sm | 1/400 | D9M8I |
| SFTPC | 153 Eu | 1/400 | EPR19839 |
| COL4 | 154 Sm | 1/200 | EPR24281-65 |
| CES1 | 155 Gd | 1/400 | EP1376Y |
| PDGFRB | 156 Gd | 1/100 | MAB1263 |
| AQP4 | 158 Gd | 1/400 | EPR24281-65 |
| CD68 | 159 Tb | 1/400 | KP1 |
| PLIN2 (perilipin2) | 160 Gd | 1/400 | 2C5H8 |
| CD20 | 161 Dy | 1/100 | 2H7 |
| KRT5 | 162Dy | 1/200 | SP27 |
| P16 | 163 Dy | 1/100 | 1D7D2A1 |
| Arginase1 | 164Dy | 1/200 | D4E3M |
| anti-rabbit 2 antibody/DCN | 166 Er | 1/200 DCN, 1/200 166Er-anti-rabbit | 16813424 |
| Ki67 | 168 Er | 1/200 | B56 |
| COL1A1 | 169Tm | 1/1000 | 3169023D |
| COL15A1 | 171 Yb | 1/100 | OTI1C5 |
| YAP1 | 173 Yb | 1/600 | NB110-58358 |
| Keratin8/18 | 174Yb | 1/200 | C51 |
| TAZ | 175 Lu | 1/600 | NB110-58359 |

**Table E1 Hyperion IMC antibody information** Antibodies used in Hyperion IMC listed in the order of name, conjugated metal, concentration used and clone information.
